# ATP-competitive and allosteric inhibitors induce differential conformational changes at the autoinhibitory interface of Akt

**DOI:** 10.1101/2022.07.14.499806

**Authors:** Alexandria L Shaw, Matthew AH Parson, Linda Truebestein, Meredith L Jenkins, Thomas A Leonard, John E Burke

## Abstract

The protein kinase Akt is a master regulator of pro-growth signalling in the cell. Akt is activated through its targeted recruitment to phosphoinositides, leading to disruption of the autoinhibitory interface between the kinase and pleckstrin homology (PH) domains. Hyper activation of Akt is common in oncogenic transformation, with multiple oncogenic activating mutants identified in Akt. This has led to the development of potent and selective ATP-competitive and allosteric inhibitors for Akt. Paradoxically, some ATP-competitive Akt inhibitors cause hyperphosphorylation of Akt. Here, using hydrogen deuterium exchange mass spectrometry (HDX-MS), we interrogated the conformational changes upon binding to the Akt active site inhibitor A-443654, and the Akt allosteric inhibitor MK-2206. We compared the conformational changes that occurred for each inhibitor under three different states of Akt: **i-**inactive monophosphorylated, **ii-**partially active tris-phosphorylated [T308, T450, S473], and **iii-**fully activated, tris-phosphorylated bound to PIP_3_ membranes. The allosteric MK-2206 inhibitor results in large scale allosteric conformational changes in all states, and restricts membrane binding through sequestration of the PH domain. Binding of the A-443654 inhibitor led to large scale allosteric conformational changes in both the monophosphorylated and phosphorylated states, leading to an alteration in the autoinhibitory PH-kinase interface. We also observed increased protection in the PH domain upon membrane binding in the presence of A-443654, suggesting that the PH domain is more accessible for membrane binding. This work provides unique insight into the autoinhibitory conformation of the PH and kinase domain and dynamic conformational changes induced by Akt inhibitors, and has important implications for the design of Akt targeted therapeutics.

## Introduction

The AGC family protein kinase Akt (also referred to as Protein Kinase B) is a serine/threonine kinase that is involved in cellular growth, proliferation, differentiation, and metabolism(1–3). There are three Akt isoforms (Akt1, Akt2, and Akt3), with all being activated downstream of the phosphoinositide 3-kinase (PI3K) signalling pathway (4). PI3K generates the lipid second messenger PIP_3_ which directly activates Akt, and recruits the Akt activator phosphoinositide dependent protein kinase 1 (PDK1)(5, 6). Activated Akt phosphorylates a number of downstream substrates, which play critical roles in a myriad of cellular signalling functions. Akt is composed of two domains, a bi-lobal protein kinase domain, and a N-terminal pleckstrin homology (PH) domain that inhibits protein kinase activity and binds to the phosphoinositides PIP_3_ and PI(3,4)P_2_, mediating Akt activation (Fig. 1A). The activity of Akt is further regulated by a plethora of post-translational modifications, including phosphorylation, methylation, SUMOylation, glycosylation, ubiquitination and acetylation(7). Aberrant activated PI3K/Akt signalling is a key driver in human cancers, with frequent activating oncogenic mutations identified in numerous members of the PI3K/Akt signalling pathway(4).

**Figure 1:**
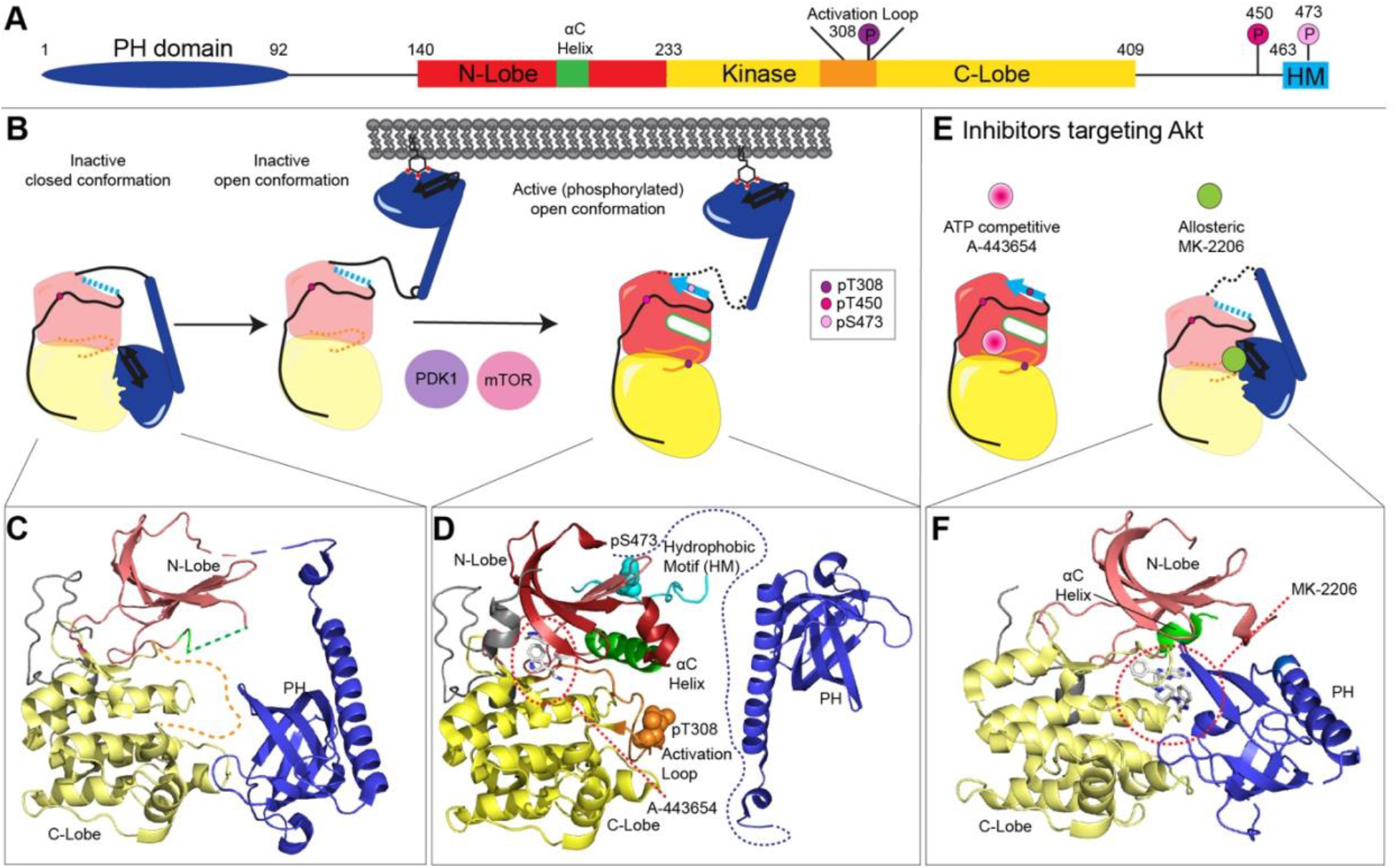
**A)** Domain schematic of Akt1 highlighting critical regions in protein kinase activation including the phosphorylation sites at T308, T450, and S473, the alpha-C helix (αC), the activation loop, and the hydrophobic motif (HM). **B)** Cartoon representation of the structural basis of Akt1 activation by PIP_3_ and phosphorylation. This model highlights the canonical activation pathway, where Akt1 binds to PIP_3_ through its PH domain, releasing the auto-inhibitory contact, and exposing the activation loop and hydrophibic motif to kinases, leading to phosphorylation of the activation loop (308) and hydrophibic motif (473). Phosphorylation of these regions leads to ordering of the αC helix (green) and activation loop, and activation of protein kinase activity. **C)** Structural model of inactive Akt1 in its closed conformation (PDB: 7APJ) (10) used to map HDX-MS data. Note that much of the N-lobe is disordered in the absence of phosphorylation, including the αC helix, as well as the activation loop in the C-lobe. **D)** Structural model of active (phosphorylated) Akt1 with disengaged PH domain bound to active site inhibitor A-443654 (white) used to map HDX-MS data. The model was generated using the structure of active Akt1 bound to ATP and peptide substrate (4EKK) (33), with the position of A-443654 overlaid from the structure of A-443654 bound to Akt2 (2JDR) (47). The PH domain is released upon membrane binding, and the orientation of the PH domain is tangential to that of the kinase domain. **E)** Inhibitors target Akt1. ATP-competitive inhibitor A-443654 is ATP competitive, with the allosteric inhibitor MK-2206 reorienting the PH domain relative to the kinase Domain. **F)** Structural model of Akt1 bound to the allosteric inhibitor MK-2206 used to map HDX-MS data (PDB: 3O96).

Akt is maintained in an inactive conformation in the absence of pro-growth signals, which is dependent on an inhibitory interaction between the PH domain and the kinase domain (8–11) (Fig. 1B, 1C). The nanobody stabilised structure of full length Akt revealed that the PH domain interacts with the C-lobe of the kinase domain, with the kinase domain binding at the PH domain phosphoinositide binding pocket (10, 12). This interface is disrupted upon binding PIP_3_ or PI(3,4)P_2_ containing membranes (6, 11, 13, 14) (Fig. 1B). Upon the autoinhibitory PH domain interface being disrupted Akt can be phosphorylated at T308 in the activation loop by PDK1 (15, 16), and at S473 in the hydrophobic motif by mTORC2 directly (17) or indirectly through phosphorylation of the Tor interacting Motif (18). Phosphorylation of T308 causes structural rearrangements of the activation loop leading to ordering of the αC helix in the N-lobe, and is essential in activating Akt kinase activity (19). Phosphorylation of S473 leads to engagement of the hydrophobic motif with the N-lobe, further stabilising the active conformation of the αC helix(19, 20) (Fig. 1B, 1D). In addition to these activating phosphorylation sites, Akt is constitutively phosphorylated at the turn motif (T450) which plays a critical role in protein stability (21). The exact role of phosphorylation in activation of Akt is controversial, with protein semi-synthesis and segmental labelling approaches (lacking stabilising turn motif phosphorylation) showing that S473 phosphorylation leads to PH domain disengagement (22, 23), and others showing >100 fold activation in kinase assays of doubly phosphorylated Akt1 upon PH domain removal (24), as well as studies using a truncated PH-kinase linker showing that the PH domain interface is maintained in T308, S473 phosphorylated Akt (11, 13).

There have been extensive efforts to develop Akt inhibitors as therapeutics due to the involvement of oncogenic activating mutations in PI3K and Akt in cancer and overgrowth disorders (25–27). The activating E17K mutation in the PH domain of Akt1 occurs in ∼2-3% of urinary and bladder cancers, and is a driver mutation in the rare Proteus overgrowth disease (25, 27). Multiple types of Akt inhibitors have been developed, including ATP-competitive active site inhibitors, allosteric inhibitors, and covalent allosteric inhibitors (Fig. 1E). Akt inhibitors are in clinical trials, including selective ATP-competitive inhibitors (AZD5363, GDC-0068, etc) and allosteric inhibitors (MK-2206, ARQ092, etc) (28). The active site inhibitor A-443654 inhibits phosphorylation of AKT’s downstream effectors, although it causes hyperphosphorylation of AKT at residues T308 and S473 *in vivo* (29, 30). This is also observed for AZD5363 (31) and GDC-0068 (32), suggesting it is a general feature of ATP-competitive inhibitors. Presence of ATP-competitive inhibitors could increase phosphorylation by making it a more suitable substrate for its upstream activators (e.g., PDK1 & mTOR) or by making it a worse substrate for phosphatases. There is data showing that ATP-competitive inhibitors or ATP protect the pT308 and pS473 phosphorylation sites from phosphatase access (33, 34). The exact conformational changes induced by inhibitor binding that mediate this process are still not fully understood at a molecular level. In contrast allosteric inhibitors do not target the conserved ATP binding site, but instead bind at an allosteric pocket between the kinase and PH domain preventing Akt phosphorylation and activation (35, 36). The structure of MK-2206 bound to Akt1 shows a distinctly different orientation of the PH domain relative to the kinase domain in the absence of the inhibitor (37) (Fig. 1C vs 1F). The altered interface of the PH domain with MK-2206 is supported by NMR (23) and FRET (8) studies showing a difference in the conformation of the PH domain in apo and MK-2206 bound Akt1.

To fully explore the conformational dynamics of Akt1 inhibition by both ATP-competitive and allosteric inhibitors we have utilised hydrogen deuterium exchange mass spectrometry (HDX-MS). Differences in conformation were analysed under three distinct states of Akt activation: the fully inactive unphosphorylated at T308 and S473 state, partially active fully phosphorylated at T308 and S473, and the fully phosphorylated PIP_3_ activated state. HDX-MS revealed extensive conformational differences in the PH-kinase interface upon binding allosteric Akt inhibitors, suggesting a unique interface. ATP-competitive inhibitors induced large scale conformational changes throughout the kinase domain and caused exposure of the PH domain in both the monophosphorylated and phosphorylated states. Overall, our findings provide novel molecular insight into the regulatory mechanisms of Akt1 regulation by the PH domain and phosphorylation, and how these are altered by small molecule inhibitors.

## Results

The conformation of Akt is strongly dependent on regulatory post-translational modifications. To determine the full complement of conformational changes induced by ATP-competitive and allosteric inhibitors we needed to examine both inactive and active Akt1. This required analysing Akt1 in both fully phosphorylated and monophosphorylated (pT450) states. We purified two states of Akt1: Akt1 DrLink (which will be referred to as Akt1 throughout the manuscript), which is monophosphorylated at the T450 (turn motif) and Akt1 DrLink 3P (referred to as Akt1 3P throughout the manuscript), which is phosphorylated at the turn motif as well as T308 (activation loop), and S473 (hydrophobic motif). Both constructs were expressed in Sf9 insect cells, with phosphorylated Akt1 3P generated by treating Sf9 cells with the inhibitors A443654 and okadaic acid during protein expression, with *in vitro* phosphorylation of T308 by PDK1 during protein purification (10). The percent phosphorylation for all sites was characterised by mass spectrometry. The Akt1 construct was ∼80% phosphorylated at T450, with no detectable additional phosphorylation, and the Akt1 3P was ∼90% phosphorylated at T450, ∼98% phosphorylated at T308, and ∼95% phosphorylated at S473 (source data). Both constructs are in the context of a truncated PH-kinase domain linker derived from the zebrafish *Danio rerio* homolog as it decreases conformational heterogeneity while maintaining equal PIP_3_ binding affinity, similar activation by PIP_3_, and Akt1 kinase activity (11) (Fig. S1A/B). Throughout this manuscript, human Akt1 numbering is used to omit confusion, however it is important to note that there is an offset of +7 residues to return to human numbering, as a result of the shortened linker, beginning at the kinase domain’s N-lobe.

### Use of hydrogen deuterium exchange mass spectrometry to analyse inhibitor binding to Akt1

We carried out HDX-MS experiments on both autoinhibited Akt1 and partially activated Akt1 3P in the presence and absence of active site (A-443654) and allosteric (MK-2206) inhibitors to define conformational changes accompanying inhibitor binding. HDX-MS measures amide hydrogen exchange with deuterated solvent, with the main determinant of amide exchange being the presence and stability of secondary structure. It acts as a readout of protein conformational dynamics, and is useful in mapping protein-protein (38), protein-membrane (11), and protein-small-molecule interactions (39). Deuterium incorporation is localized at peptide-level resolution through the generation of pepsin-generated peptides. The HDX-MS coverage of Akt1 was composed of 147 identified peptides that spanned 98.9% of the primary sequence of Akt1, including multiple charge states for a total of 251 monitored isotopic peaks, and the HDX-MS coverage of Akt1 3P was composed of 143 identified peptides that spanned 98.9% including multiple charge states for a total of 166 monitored isotopic peaks. HDX-MS experiments compared the dynamics of Akt1 and Akt1 3P with the inhibitors A-443654 and MK-2206. The full raw deuterium incorporation data for all HDX-MS experiments are provided in the Source data.

### HDX-MS reveals conformational changes induced by active site inhibitor A-443654 in autoinhibited Akt1

Initial HDX-MS experiments were carried out with autoinhibited Akt1 in the presence of the active site inhibitor of A-443654. Experiments were carried out at saturating levels of A-443654 (final concentration of 1.5 µM), with deuterium incorporation measured over five time points of exchange (3, 30, 300, 3000 seconds at 18°C and 3 seconds at 1°C, which is referred to as 0.3 sec in all graphs and the source data).

Multiple regions of Akt1 had significant alterations in deuterium incorporation upon incubation with A-443654 (significant change in exchange defined as greater than both 4.5% and 0.45 Da at any time point in any peptide with an unpaired two-tailed t-test p value <0.01). These included significant decreases in exchange in the kinase domain, and significant increases in exchange in the PH domain (Fig. 2A, 2C, 2F, S2A, S3A). All regions in the N-lobe and C-lobe in contact with the inhibitor in the crystal structure showed significantly decreased exchange, with one of the largest % differences in exchange occurring at the hinge region between the two lobes (229-235). There were also large decreases in exchange in the region spanning the αC helix (176-198, 198-210), which is unstructured in the inactive configuration of Akt1in the absence of nucleotides. There were also decreases in regions of the N-lobe that were unstructured in the inactive Akt1 crystal structure in the absence of the C-terminal tail, suggesting that inhibitor binding stabilizes the N-lobe in inactive Akt1. Extensive allosteric conformational changes were observed in the C-lobe, with large decreases in exchange observed at the C-terminal end of the C-lobe, near the PH domain interface. Intriguingly, the largest % difference in exchange was located from 358-365, which is greater than 20 Å from the inhibitor binding site. Together, this suggests that ATP-competitive inhibitors reshape the conformation of the PH domain interface in the kinase domain. There was a small increase in exchange in the PH domain spanning residues 76-95. The rest of the PH domain showed similar exchange. Previous experiments examining oncogenic mutations (D323A&D325A) in the activation loop of Akt1 that led to disengagement of the PH-kinase interfaces showed much more extensive increases in exchange (Fig. S3D) (11), suggesting that there is not complete disengagement of the PH domain upon binding to ATP-competitive inhibitors. Overall, this data suggests allosteric conformational changes in the C-lobe induced upon A-443654 binding, leading to a weakening of the PH-kinase interface.

**Figure 2.**
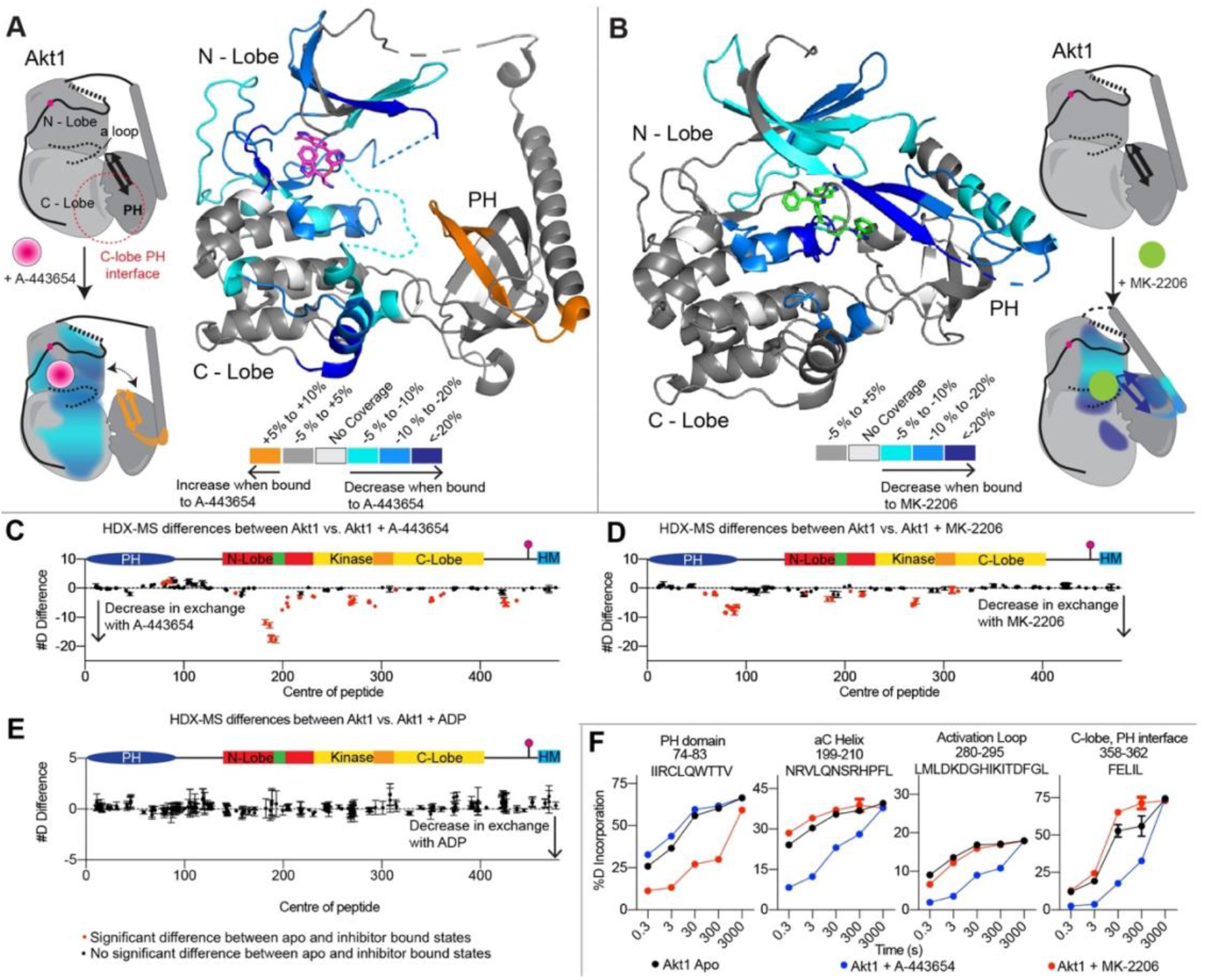
HDX-MS differences upon inactive Akt1 binding A-443654 and MK-2206. **A+B**. Peptides showing significant differences in deuterium exchange (defined as >4.5%, 0.45 Da, and p<0.01 in an unpaired two-tailed T-test at any timepoint) upon addition of A-443654 (**A**) or MK-2206 (**B**). Differences in panel A are mapped on the structural model from Fig 1C (7APJ), and differences in panel B are mapped on the structural model from 1F (3O96). The difference is indicated by the legend, with a cartoon model highlighting the location of these significant differences. **C-E**. Sum of the number of deuteron difference of Akt1 upon binding 1.5 μM A-443654 **(C)**, 1.5 μM MK-2206 **(D)**, and 1 mM ADP **(E)**, analysed over the entire deuterium exchange time course for Akt1. Each point is representative of the centre residue of an individual peptide. Peptides that met the significance criteria described in A are coloured red. Error is shown as standard deviation (n=3). **F**. Selected deuterium exchange time courses of Akt1 peptides that showed significant decreases and increases in exchange. Error is shown as standard deviation (n=3).

To determine which of these changes were caused by occupancy of the active site by substrate/product, we carried out HDX-MS experiments with 1 mM ADP, which is a concentration that should saturate the active site. No significant differences in exchange were observed (Fig. 2E), showing a clear difference between ATP-competitive inhibitors.

### HDX-MS reveals conformational changes induced by allosteric inhibitor MK-2206 in autoinhibited Akt1

To compare conformational changes upon allosteric inhibitor binding, we carried out HDX-MS experiments with the clinical allosteric inhibitor MK-2206. HDX-MS experiments were carried out at saturating levels of MK-2206 (final concentration of 1.5 µM), with deuterium incorporation measured over five time points of exchange (3, 30, 300, 3000 seconds at 18°C and 3 seconds at 1°C, which is referred to as 0.3 sec in all graphs and the source data).

There were multiple regions of significantly decreased deuterium exchange in the kinase and PH domains (Fig. 2B, 2D, 2E). There was decreased exchange in the PH domain spanning a region from 57-104, with the largest % difference occurring from residues 74-88 (PIP_3_ binding motif), with this region being in contact with MK-2206 in the corresponding crystal structure (37) (Fig. 2B/D/E). Decreases in exchange in the kinase domain were observed (Fig. 2B/D/E) spanning residues 196-198, 213-225, 259-274, and 298-321. The structure of inactive Akt1 bound to MK-2206 (PDB 3O96) showed a reorientation of the PH domain compared to the inactive structure (Fig. S4C), where MK-2206 sequesters the PIP_3_ binding motif through the formation of a unique PH-kinase contact site, mediated by MK-2206 binding (Fig. 1F). Our HDX-MS data is consistent with this altered orientation of the PH domain.

### HDX-MS reveals conformational changes induced by active site inhibitor A-443654 in phosphorylated partially active Akt1

To compare possible unique conformational changes upon A-443654 binding to partially activated Akt1 3P, we carried out HDX-MS experiments with Akt 3P. Our observation that ATP-competitive inhibitors weakened the auto-inhibitory kinase-PH interface also allowed us to interrogate the potential role of phosphorylation in disrupting this interface.

There were multiple regions of significantly decreased deuterium exchange in the kinase domain, as well as a significant increase in exchange in the PH domain (Fig. 3A/C/E). Similar to before there were significant decreases in regions directly in contact with the inhibitor, the αC helix, and part of the C-lobe PH domain interface. There were also similar increases in exchange in the PH domain (74-95), consistent with what was observed in inactive Akt1. Intriguingly there was no significant difference observed at the 358-365 region at the C-lobe-PH interface (Fig. 2F/3E). Overall, outside of the difference at region 358-365, there were similar decreases in exchange observed in both Akt1 and Akt1 3P upon inhibitor binding, with the main difference being the intensity of the difference (Fig. 2C vs Fig 3C). Both experiments were carried out under saturating concentrations of inhibitor binding, so this difference reflects intrinsic conformational differences. The larger differences in exchange observed in the Akt1 sample are likely due to the increased protection of the unstructured regions in the N-lobe. Critically, in both samples there was increased exchange in the PH domain upon A-443654 binding, suggesting weakening of the PH-kinase domain interface. This result also provides additional evidence that the PH-kinase domain autoinhibitory interface is maintained in phosphorylated Akt1.

**Figure 3.**
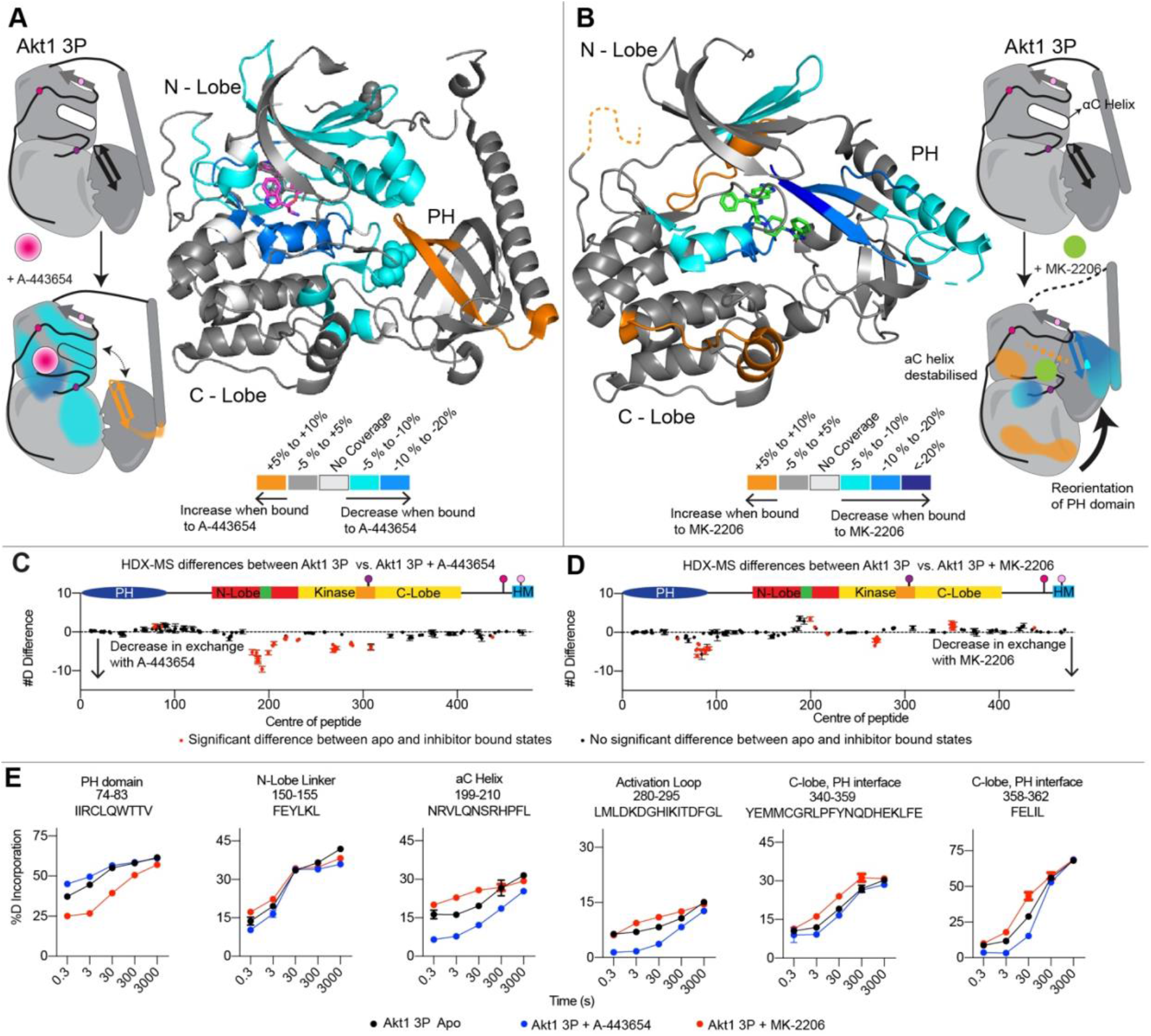
HDX-MS differences upon partially active phosphorylated Akt1 3P binding A-443654 and MK-2206. **A+B**. Peptides showing significant differences in deuterium exchange (defined as >4.5%, 0.45 Da, and p<0.01 in an unpaired two-tailed T-test at any timepoint) upon addition of A-443654 (**A**) or MK-2206 (**B**). Differences in panel A are mapped on the structural model from Fig 1D (4EKK), with the PH domain oriented from 7APJ and the T308 phospho site shown as spheres, and differences in panel B are mapped on the structural model from 1F (3O96). The difference is indicated by the legend, with a cartoon model highlighting the location of these significant differences. **C+D**. Sum of the number of deuteron difference of Akt1 3P upon binding 2.2 μM A-443654 or, **(C)** 2.2 μM MK-2206 **(D)**, analysed over the entire deuterium exchange time course for Akt1 3P. Each point is representative of the centre residue of an individual peptide. Peptides that met the significance criteria described in A are coloured red. Error is shown as standard deviation (n=3). **E**. Selected deuterium exchange time courses of Akt1 3P peptides that showed significant decreases and increases in exchange upon inhibitor binding. Error is shown as standard deviation (n=3).

### HDX-MS reveals that MK-2206 restructures and repositions αC helix and C-lobe:PH domain interface residues

Coincubation of Akt1 3P with MK-2206 revealed multiple regions of significantly decreased and increased deuterium exchange in the kinase and PH domain (Fig. 3B/D). Decreases in deuterium were observed throughout the PH domain (Fig. 3B/D/E). In the kinase domain, decreases in deuterium exchange occurred at residues 213-225 and 259-280 with residues 264-274 having a moderate decrease (Fig. 3B/D/E). Additionally, small increases in deuterium exchange were also observed in the kinase domain, specifically at the αC helix (residues 191-210) and C-lobe (residues 342-359) near the C-lobe:PH interface (Fig. 3B/D/E). This is consistent with MK-2206 binding being incompatible with formation of the αC helix due to steric clashes between MK-2206, and the active conformation of the αC helix (Fig. S4C). The increased exchange seen at the C-lobe:PH domain interface is also consistent with a rotation of the PH domain as predicted by the crystal structure (Fig. S1F).

### HDX-MS analysis of fully activated membrane bound Akt1 bound to active site and allosteric inhibitors

To fully define the role of inhibitors in modifying the PH-kinase domain interface, we required a system to probe conformational changes that occurred in a fully activated Akt1 with the autoinhibitory PH interface disrupted. To do this we carried out HDX-MS experiments on Akt1 pre-incubated to lipid vesicles composed of phosphatidylethanolamine (PE), phosphatidylserine (PS), cholesterol, phosphatidylcholine, and phosphatidylinositol 3,4,5-trisphosphate (PIP_3_) in the presence and absence of inhibitors A-443654 and MK-2206. We have previously shown that binding to PIP_3_ vesicles leads to release of the PH domain for both inactive and phosphorylated Akt1 (10, 11). An important note on these experiments is that HDX-MS changes report on the full set of conformational changes that occur, which may include altered association with membranes due to inhibitors, as well as the inherent conformational changes that occur due to inhibitor binding.

There were multiple regions of significantly decreased deuterium exchange for A-443654 binding to Akt1 3P pre-associated with 5% PIP_3_ vesicles, including in both the kinase and PH domains (Fig. 4A/C). Intriguingly there were no increases in exchange observed in the PH domain. Decreases in exchange in the kinase domain were similar to those observed in the absence of membranes, occurring in regions encompassing the αC helix, the ATP binding pocket, as well as changes covering the activation loop and C-lobe:PH interface (Fig. 4A/C/G). In the PH domain, when bound to 5% PIP_3_ Vesicles in the presence of A-443654 there is a moderate decrease in deuterium exchange at residues 76-88 at the membrane binding site. Potentially this could indicate that the reorientation of the PH domain we observed in the non-membrane bound state could promote additional membrane association, with this decrease indicating enhanced Akt1 membrane binding upon inhibitor binding (Fig. 4E).

**Figure 4.**
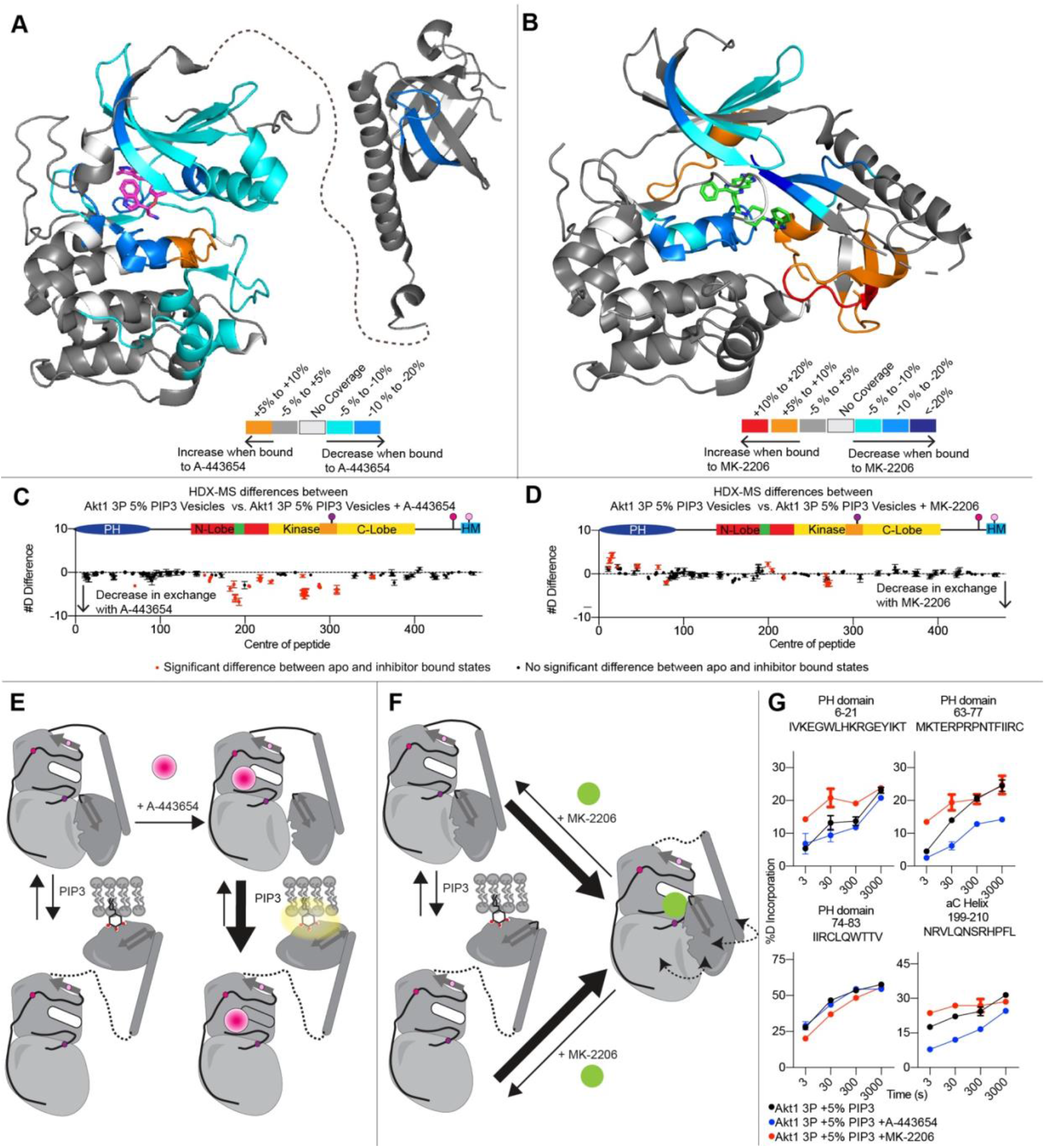
HDX-MS differences upon fully active PIP_3_ membrane bound phosphorylated Akt1 3P binding A-443654 and MK-2206. **A+B**. Peptides showing significant differences in deuterium exchange (defined as >4.5%, 0.45 Da, and p<0.01 in an unpaired two-tailed T-test at any timepoint) upon addition of A-443654 (**A**) or MK-2206 (**B**). Differences in panel A are mapped on the structural model from Fig 1D (4EKK), and differences in panel B are mapped on the structural model from 1F (3O96). The difference is indicated by the legend. **C+D**. Sum of the number of deuteron difference of PIP_3_ membrane bound Akt1 3P upon binding 2.2 μM A-443654 or **(C)** 2.2 μM MK-2206 **(D)**, analysed over the entire deuterium exchange time course for Akt1 3P. Each point is representative of the centre residue of an individual peptide. Peptides that met the significance criteria described in A are coloured red. Error is shown as standard deviation (n=3). **E+F**. Cartoon model indicating altered membrane binding of Akt1 upon binding either A-443654 (**E**) or MK-2206 (**F**). HDX-MS results are consistent with enhanced membrane binding in the presence of A-443654, and decreased membrane binding in the presence of MK-2206, mediated by enhanced PH domain disengagement in A-443654 and decreased PH domain disengagement in MK-2206. **E**. Selected deuterium exchange time courses for peptides of Akt1 3P in the presence of 5% PIP_3_ vesicles that showed significant decreases and increases in exchange upon inhibitor binding. Error is shown as standard deviation (n=3).

Experiments examining HDX-MS differences upon MK-2206 binding to Akt1 3P in the presence of 5% PIP_3_ vesicles revealed multiple regions of significantly decreased and increased deuterium exchange in the kinase domain and PH domain (Fig. 4B/D/G). Decreased deuterium exchange occurred at the interface of the PH domain with MK-2206 at residues 76-85, and increased exchange at residues 6-26, 38-55, and 65-75 (Fig. 4B/D/G). These changes in deuterium uptake might suggest that MK-2206 outcompetes PIP_3_ binding, essentially removing the PH domain from the plasma membrane and sequestering the PIP_3_ binding site (Fig. 4B/F), preventing membrane association. This is consistent with our previous results showing greatly reduced membrane binding when allosteric inhibitors are present (10). In the kinase domain, decreases in deuterium exchange occurred at residues 213-225 and 259-280 (Fig. 4B/D/G). Additionally, a small increase in deuterium exchange was also observed in the kinase domain, similar to the membrane free Akt1 3P +MK-2206 at the αC helix (191-210) (Fig. 4B/D/G). The observed increase in deuterium exchange at the C-lobe (residues 342-359) near the C-lobe:PH interface shown when comparing Akt1 3P apo to Akt1 3P +MK-2206 was not seen in the presence of PIP_3_. This is potentially due to this region being already exposed due to the release of the PH domain upon binding PIP_3_ membranes.

## Discussion

The PI3K/Akt pathway is the most frequently activated pathway in human cancer, with mutations accounting for >25% of all cancers (40). This has driven intense interest in developing inhibitors for both the lipid kinase PI3Ks and Akt. Akt inhibitors have advanced through pre-clinical and clinical trials, with the MK-2206 allosteric inhibitor and the ATP-competitive inhibitor GSK2141795 in phase II clinical trials (41). Partially limiting the application of Akt inhibitors has been the observation of hyperphosphorylation of Akt1 upon binding ATP-competitive inhibitors (29), including the clinical candidate GSK2141795 (42). Inhibitor mediated hyperphosphorylation is proposed to be mediated through protection from dephosphorylation (33, 34), although the full molecular mechanism underlying this phenomenon is not completely understood. Critical to the development of Akt inhibitors is understanding the conformational changes underlying inhibitor binding, as well as the conformational changes that underly Akt activation. Here we describe the conformational changes occurring with both active site and allosteric inhibitors across the full conformational landscape of Akt activation.

Different active site ATP-competitive inhibitors all lead to hyperphosphorylation of Akt1 upon inhibitor treatment. We observed that A-443654 caused decreased exchange throughout the kinase domain, and increased exchange in the PH domain, which was conserved in both monophosphorylated and phosphorylated Akt1. This increased exchange in the PH domain in both Akt1 and Akt1 3P, which is indicative of a weakening of the inhibitory interface is consistent with our previous data showing that the PH-kinase interface is conserved in both inactive and phosphorylated Akt1 (10, 11). Intriguingly, the increased exchange observed is much smaller than that observed for oncogenic mutants at the PH-kinase interface (11), highlighting that the interface is not completely disrupted, and is only weakened. An unanswered question from our previous structural data on inactive Akt1 was how does the PH domain interface remain intact, as the crystal structure of phosphorylated Akt1 showed an orientation of the activation loop that would clash with the PH domain orientation from inactive Akt1 (10, 19, 20). Initial insight into this difference is potentially provided by the difference in A-443654 binding between the inactive monophosphorylated Akt1 and partially active Akt1 3P. In inactive Akt1 there is a large decrease in exchange in the C-lobe of the kinase domain at the interface with the PH domain upon A-443654 binding. This decrease was not observed in the Akt 3P construct upon A-443654 binding. The PH domain has the same increase, suggesting that there is a shared weakening of the autoinhibitory interface with the PH domain, but suggests that the PH domain might form an altered interface with the C-lobe in phosphorylated Akt1. Further structural and biochemical data will be required to define the exact molecular basis of the PH-kinase interface in phosphorylated Akt1.

The crystal structure of unphosphorylated Akt1 in complex with MK-2206 showed an altered orientation of the PH domain compared to inactive Akt1 (10, 37), where the PH domain forms a more extensive interface with the kinase domain, with the inhibitor binding between both the PH and kinase domains. The PH domain rotates towards the position of the location of the αC helix in phosphorylated Akt1, sequestering its PIP_3_ binding motif. NMR studies examining the interaction of allosteric inhibitors have also observed data consistent with an altered orientation of the PH domain compared to inactive Akt1 (23). We observed differences in exchange in Akt1 and Akt1 3P consistent with altered packing of the PH domain. The primary difference between Akt1 and Akt1 3P upon MK-2206, was the extensive increases in exchange observed in the kinase domain upon Akt1 3P, with the majority of these changes not observed in Akt1. When Akt1 3P is bound to MK-2206, there was a large increase in exchange in the αC helix, consistent with a disruption of the active site. This increase was not seen for MK-2206 binding in inactive Akt1, although the region spanning the αC helix is much more dynamics in Akt1, consistent with this helix not forming in inactive Akt1 (10). There were also increases in exchange in the C-lobe in Akt1 3P upon MK-2206 binding. This is likely also due to difference in the orientation of the PH domain for Akt1 3P compared to Akt1.

To validate the weakening of the PH domain observed for both Akt1 and Akt1 3P required testing a condition where the PH domain is likely to be fully disengaged. HDX-MS comparison between Akt1 3P alone and bound to PIP_3_ membrane vesicles showed increased exchange in the kinase domain, and decreased exchange in the PH domain (10), consistent with disengagement of the PH-kinase interface. HDX-MS experiments carried out with A-443654 for membrane bound Akt1 showed similar changes in the kinase domain, however, there was now no longer increased exchange in the PH domain. Intriguingly we observed decreased exchange in a portion of the PH domain. We propose that this likely due to enhanced membrane binding of Akt1 in the presence of ATP-competitive inhibitors due to weakened autoinhibitory interface, and enhanced exposure of the PIP_3_ binding site in the PH domain. Experiments with MK-2206 showed an opposite trend, with there now being increased exchange in multiple regions of the PH domain, indicative of decreased membrane binding. This is consistent with our previous membrane binding experiments showing decreased membrane recruitment with allosteric Akt1 inhibitors (10).

Overall, our study has revealed unique insight into conformational changes underlying Akt1 activation, as well as distinct conformational differences between activation states for both active site and allosteric inhibitors. We identified conformational changes that are distinct between both inhibitor types and activation states. Further exploration of the dynamic regulation of Akt1 by oncogenic mutations and other post-translational modifications, along with how they can be targeted by ATP-competitive and allosteric inhibitors could reveal unique approaches for Akt1 therapeutic development.

## Acknowledgements

JEB is supported by an operating grant from the Cancer Research Society (CRS-843232), with salary support from the Michael Smith Foundation for Health Research Scholar award (MSFHR-17686). TAL was supported by Austrian Science Fund Grants P30584 and P33066, and LT was supported by the Hertha Firnberg Postdoctoral Fellowship T915.

## Methods

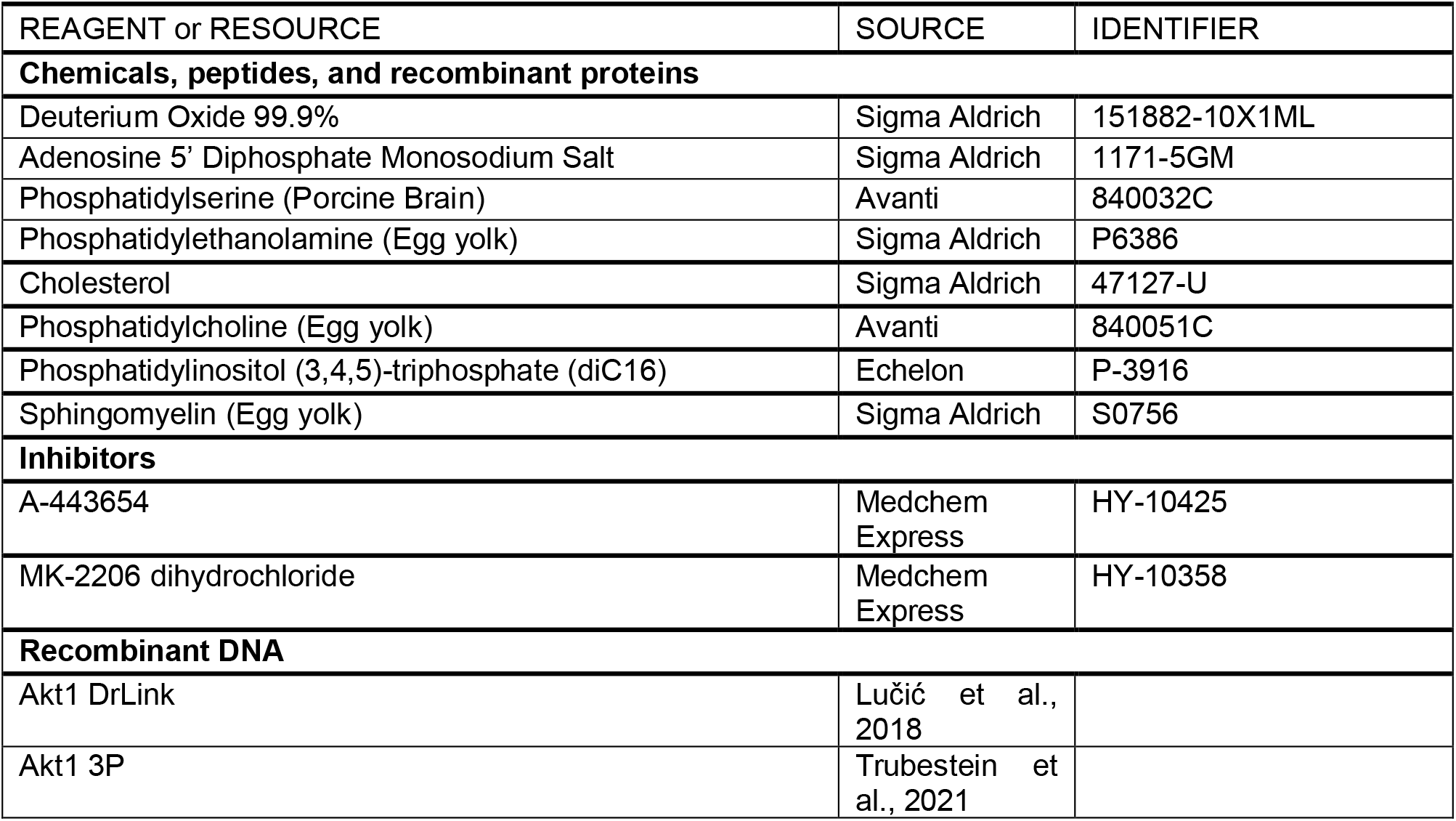
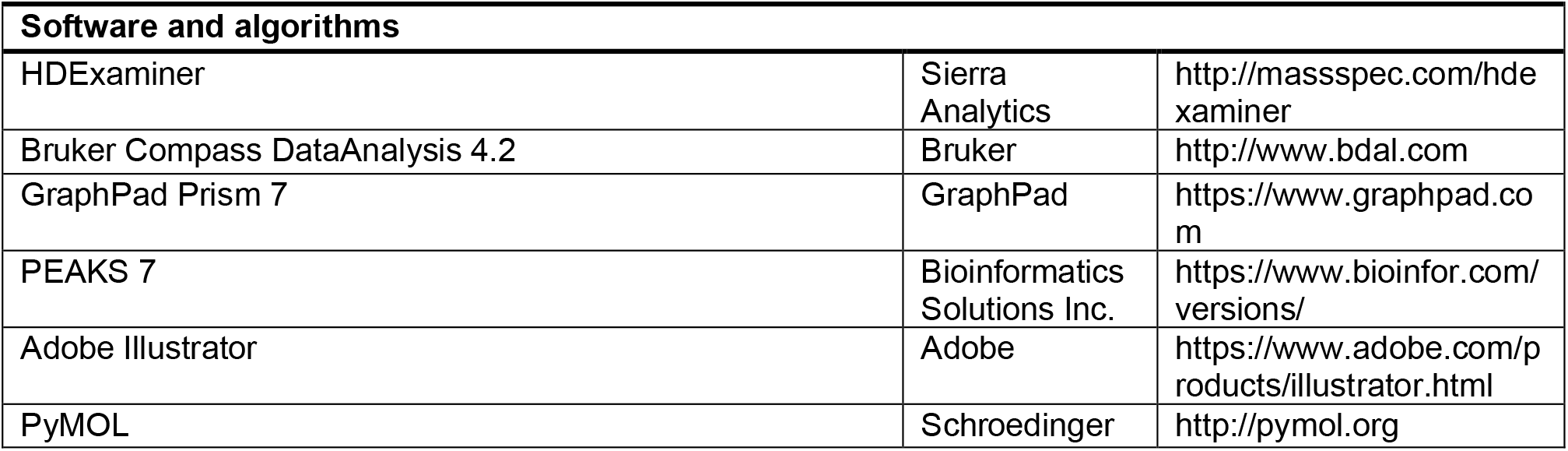

### Protein Purification

Akt1 was expressed and purified according to the protocol for Akt1_WT_ previously reported (11). Akt1 3P phosphorylated on T308, T450 and S473 was prepared as previously described (10). Briefly, Sf9 cells expressing Akt1 (Akt1) or co-expressing Akt1 S475A T479R and human PDK1 (Akt1 3P) for 72 h, with Akt1 3P samples having the Akt inhibitor A-443654 added at 1 µM for 18 h and okadaic acid (100 nM) for 60 min prior to harvesting. Both Akt1 samples were purified by resuspending Sf9 cells in lysis buffer (50 mM Tris, pH 8, 100 mM NaCl, 50 mM NaF, 1 mM TCEP (tris carboxyethylphosphine), 20mM imidazole, with protease inhibitor cocktail 2 containing 1 mM PMSF, 100 µM bestatin, 14 µM E-64, 10 µM pepstatin A, and 1 µM phosphoramidon, for 1 hr. The lysate was centrifuged at 39 000 × g for 40 min and the supernatant loaded onto StrepTrap column (GE Healthcare) in lysis buffer containing 50 mM Tris, pH 7.4, 100 mM NaCl, 50 mM NaF, 1 mM sodium orthovanadate, 10 mM b-glycerophosphate, 2 mM sodium pyrophosphate, 2 mM benzamidine, 1 mM EDTA, 1 mM TCEP and protease inhibitor cocktail containing 1 mM PMSF, 100 µM bestatin, 14 µM E-64, 10 µM pepstatin A, 1 µM phosphoramidon. The protein was washed with buffer StrepA (50 mM Tris, pH 7.4, 100 mM NaCl, 50 mM NaF, 1 mM EDTA, 1 mM TCEP) and eluted in buffer StrepB (StrepA with addition of 2.5 mM D-desthiobiotin). Peak fractions were pooled and subjected to overnight cleavage with TEV protease and further purification by high-resolution MonoQ anion exchange chromatography and size exclusion chromatography was performed in 20 mM Tris, pH 8.0, 100 mM NaCl, 50 mM NaF, 1 mM TCEP and 1 mM EDTA.

Phosphorylation of Akt1 and Akt1 3P were validated by LC mass spectrometry, with raw intensities for the phosphorylated and non-phosphorylated peptides used to determine the ratio of phosphorylation, with raw data included in the source data.

### Lipid Vesicle Preparation

Lipid vesicles containing 20% cholesterol (Sigma:47127-U), 25% egg-yolk phosphatidylcholine (PC) (Avanti: 840051C), 15% brain phosphatidylserine (PS) (Avanti: 840032C), 35% egg-yolk phosphatidylethanolamine (PE) (Sigma: P6386), and 5% phosphatidylinositol 3,4,5-trisphosphate (PIP_3_) (Echelon: P-3916) were prepared by combining lipid components dissolved in organic solvent, vigorously mixing, and evaporating the solvent under a stream of argon gas to produce an even lipid film layer. The lipid film was desiccated under a vacuum for 45 minutes and resuspended at 4 mM in lipid buffer (20mM HEPES pH 7.5, 100mM KCl) by vortexing. The resuspended lipid vesicle solutions were sonicated for 10 minutes, subjected to ten freeze-thaw cycles, and sonicated again for 5 minutes. The vesicles were then snap-frozen in liquid nitrogen and stored at −80°C.

### HDX-MS sample preparation

HDX reactions comparing Akt1 apo to Akt1 +A-443654, MK-2206, or ADP were carried out in a 20 µl reaction volume containing 15 pmol of protein and a final inhibitor concentration of 1.5 µM and final ADP concentration of 1 mM. The exchange reactions were initiated by the addition of 16 µL of D_2_O buffer (20 mM HEPES pH 7.5, 100 mM NaCl, 2 mM MgCl_2,_ 93.9% D_2_O (V/V)) to 4 µL of protein (final D_2_O concentration of 75.12%). Reactions proceeded for 3s, 30s, 300s, and 3000s at room temperature or 0.3s on ice, before being quenched with ice cold acidic quench buffer, resulting in a final concentration of 0.6M guanidine HCl and 0.9% formic acid post quench. HDX reactions comparing Akt1 3P and Akt1 3P + A-443654 or MK-2206 were carried out in 20 µL reactions containing 8.9 pmol of protein and a final inhibitor concentration of 2.2 µM. Reactions were initiated by adding 14 µL D_2_O buffer (10 mM HEPES pH 7.5, 100mM NaCl, 94.2% D_2_O) to a mixture of 4.45 µL AKT1 3P ± inhibitor and 1.55 µL of lipid buffer (20mM HEPES pH 7.5, 100mM KCl) to give a final concentration of 66% D_2_O. Exchange proceeded for 3s, 30s, 300s, and 3000s at room temperature or 0.3s on ice, before being quenched with ice cold acidic quench buffer, resulting in a final concentration of 0.6M guanidine HCl and 0.9% formic acid post quench. HDX reactions comparing Akt1 3P + PIP_3_ to Akt1 3P + PIP_3_ with inhibitors were carried out in 20µL reaction volumes with a final Akt1 3P concentration of 8 pmol. Three conditions were tested: Akt1 3P with 5% PIP_3_ vesicles (20% cholesterol, 25% PC, 15% PS, 35% PE, 5% PIP_3_), Akt1 3P with 5% PIP_3_ vesicles and 2 mM final A-443654, and Akt1 3P with 5% PIP_3_ vesicles and 2 mM final MK-2206. The final lipid vesicle concentration for all conditions was 400 µM. Prior to the exchange reactions, 4 µL of protein was incubated with 0.45 µL of lipid vesicles for 2 minutes at room temperature. HDX reactions were initiated by the addition of 1.55 µL lipid buffer and 14 µL of D_2_O buffer solution (20mM HEPES pH 7.5, 100mM NaCl, 94.2% D_2_O) to 4.45 µL of the protein-lipid solution, with a final D_2_O concentration of 66%. Exchange was carried out for four time points (3s, 30s, 300s and 3000s at room temperature). All conditions and timepoints were created and run in independent triplicate. Samples were flash frozen immediately after quenching and stored at −80°C until injected onto the ultra-performance liquid chromatography (UPLC) system for proteolytic cleavage, peptide separation, and injection onto a QTOF for mass analysis, described below.

### Protein digestion and MS/MS data collection

Protein samples were rapidly thawed and injected onto an integrated fluidics system containing a HDx-3 PAL liquid handling robot and climate-controlled (2°C) chromatography system (LEAP Technologies), a Dionex Ultimate 3000 UHPLC system, as well as an Impact HD QTOF Mass spectrometer (Bruker). The full details of the automated LC system are described in (43). The protein was run over one immobilized pepsin column (Trajan; ProDx protease column, 2.1 mm x 30 mm PDX.PP01-F32) at 200 µL/min for 3 minutes at 8°C. The resulting peptides were collected and desalted on a C18 trap column (Acquity UPLC BEH C18 1.7mm column (2.1 × 5 mm); Waters 186003975). The trap was subsequently eluted in line with an ACQUITY 1.7 μm particle, 100 × 1 mm_2_ C18 UPLC column (Waters), using a gradient of 3-35% B (Buffer A 0.1% formic acid; Buffer B 100% acetonitrile) over 11 minutes immediately followed by a gradient of 35-80% over 5 minutes. Mass spectrometry experiments acquired over a mass range from 150 to 2200 m/z using an electrospray ionization source operated at a temperature of 200C and a spray voltage of 4.5 kV.

### Peptide identification

Peptides were identified from the non-deuterated samples of Akt1 or Akt1 3P using data-dependent acquisition following tandem MS/MS experiments (0.5 s precursor scan from 150-2000 m/z; twelve 0.25 s fragment scans from 150-2000 m/z). MS/MS datasets were analysed using PEAKS7 (PEAKS), and peptide identification was carried out by using a false discovery based approach, with a threshold set to 0.1% using a database of purified proteins and known contaminants (44). The search parameters were set with a precursor tolerance of 20 ppm, fragment mass error 0.02 Da, charge states from 1-8, with a −10logP score of 26.2 and 25.8 for Akt1 or Akt1 3P respectively.

### Mass Analysis of Peptide Centroids and Measurement of Deuterium Incorporation

HD-Examiner Software (Sierra Analytics) was used to automatically calculate the level of deuterium incorporation into each peptide. All peptides were manually inspected for correct charge state, correct retention time, appropriate selection of isotopic distribution, etc. Deuteration levels were calculated using the centroid of the experimental isotope clusters. Results are presented as relative levels of deuterium incorporation and the only control for back exchange was the level of deuterium present in the buffer (74.5% and 66%). Differences in exchange in a peptide were considered significant if they met all three of the following criteria: ≥4.5% change in exchange, ≥0.45 Da difference in exchange, and a p value <0.01 using a two tailed student t-test. The raw HDX data are shown in two different formats. The raw peptide deuterium incorporation graphs for a selection of peptides with significant differences are shown in Figure S3, with the raw data for all analysed peptides in the source data. To allow for visualization of differences across all peptides, we utilized number of deuteron difference (#D) plots (Fig. 2C/D/E, 3C/D, 4C/D). These plots show the total difference in deuterium incorporation over the entire H/D exchange time course, with each point indicating a single peptide. Samples were only compared within a single experiment and were never compared to experiments completed at a different time with a different final D_2_O level. The data analysis statistics for all HDX-MS experiments are in Supplemental Table 1+2 according to the guidelines of (45). The mass spectrometry proteomics data have been deposited to the ProteomeXchange Consortium via the PRIDE partner repository (46) with the dataset identifier PXD032929.

## Supplemental Figures and Tables

**Figure S1:**
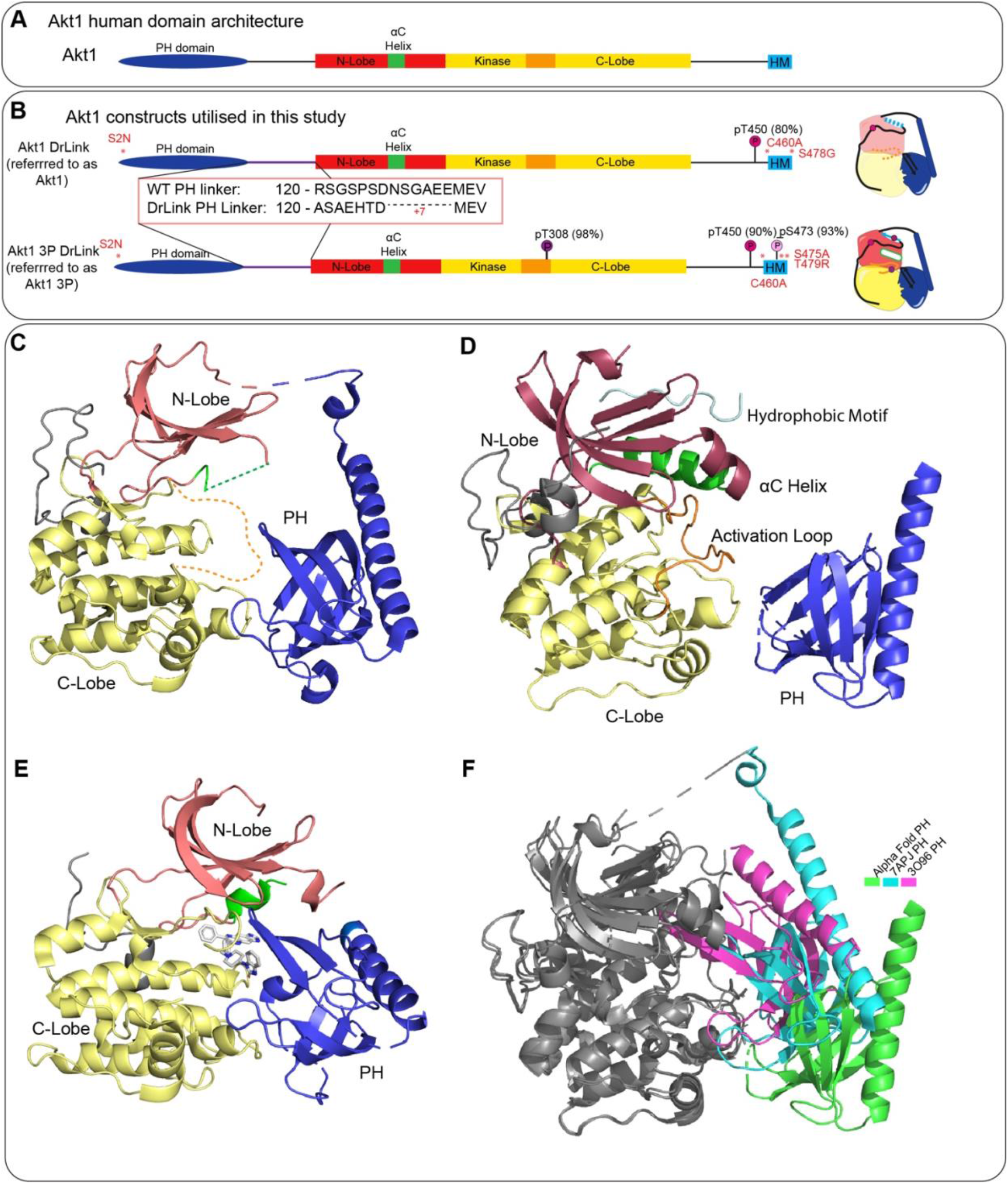
Akt1 constructs utilised in this study, and comparison of Akt1 structural models. **A**. Human Akt1 domain schematic **B**. DrLink Akt1 constructs utilized in this study, highlighting the PH-linker length used, phosphorylation sites (and their frequency), and engineered mutations. All structural models are coloured according to this schematic. **C**. Structure of inactive unphosphorylated Akt1 (PDB: 7APJ). Domains coloured per panel A/B. **D**. Predicted AlphaFold structure of inactive Akt1 (AF-P31749-F1), with regions with a pLDDT score less than 60 removed. **E**. Structure of Akt1 bound to MK-2206 (PDB: 3O96). **F**. Comparison of the PH domain positions predicted by AlphaFold, and in the structures of inactive Akt1, and Akt1 bound to the allosteric MK-2206 inhibitor.

**Figure S2:**
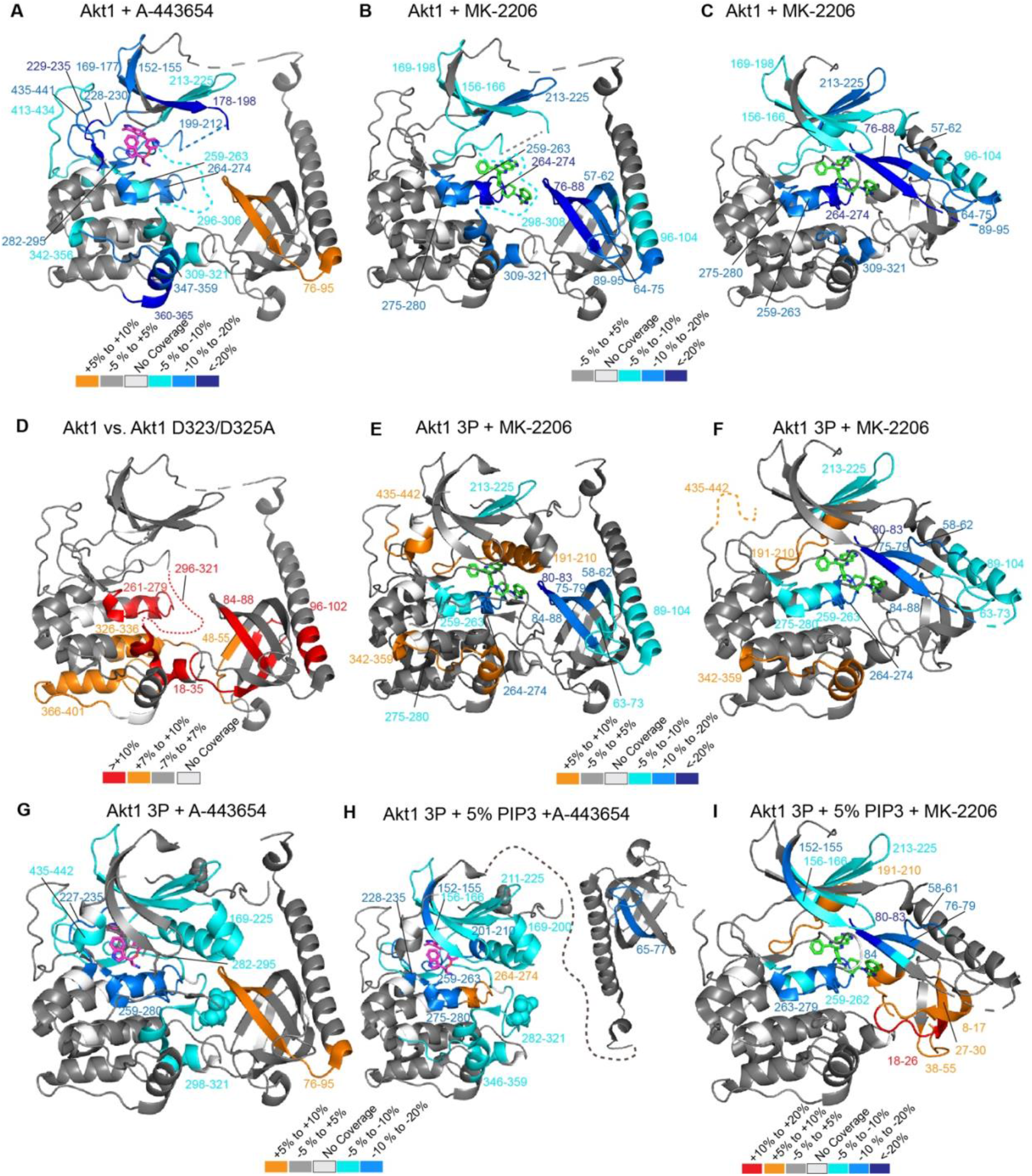
Annotated HDX-MS changes for all experiments in this manuscript. Peptides showing significant deuterium exchange differences (>4.5%, 0.45 Da, and p<0.01 in an unpaired two-tailed T-test) between various conditions. All inhibitor containing structures/models were aligned with PDB: 2JDR to show A-443654 binding and PDB: 3O96 to show MK-2206 binding locations. **(A)** Akt1 apo vs. Akt1 + A-443654 mapped to unphosphorylated Akt1(PDB: 7APJ). **(B/C)** Akt1 apo vs. Akt1 +MK-2206 mapped to **(B)** unphosphorylated Akt1 (PDB: 7APJ) and **(C)** Akt1 in complex with MK-2206 (PDB: 3O96). **(D)** Akt1 vs. Akt1 D323A/D325A mapped to unphosphorylated Akt1(PDB: 7APJ) (>7%, 0.4 Da, and p<0.01in a Student T-test). **(E/F)** Akt1 3P vs. Akt1 3P + MK-2206 mapped to **(E)** model of fully phosphorylated, PH-closed Akt1 (PDB: 4EKK (KD), 7APJ (PH)), and **(F)** Akt1 in complex with MK-2206 (PDB: 3O96). **(G)** Akt1 3P apo vs. Akt1 3P + A-443654 mapped to model of fully phosphorylated, PH-closed Akt1 (PDB: 4EKK (KD), 7APJ (PH)). **(H/I)** Akt1 3P + 5% PIP_3_ vs. **(H)** Akt1 3P + 5% PIP_3_ + A-443654 and (**I)** Akt1 3P + 5% PIP_3_ + MK-2206 mapped to **(H)** a model of fully phosphorylated, PH-closed Akt1 (PDB: 4EKK (KD), 7APJ (PH)), and **(I)** a structure of Akt1 in complex with MK-2206 (PDB:3O96). Significant differences in deuterium exchange are mapped according to the corresponding legends.

**Figure S3:**
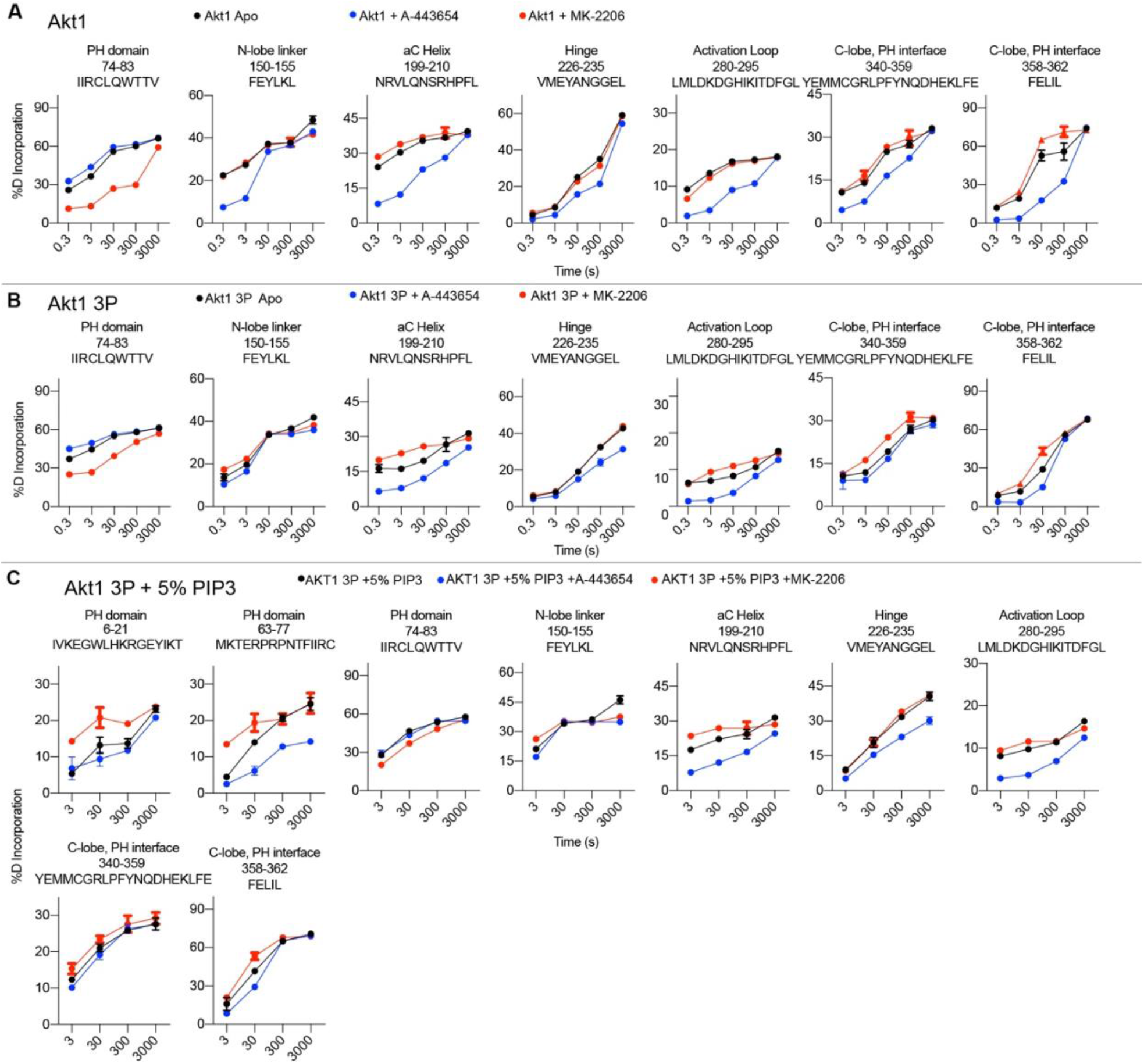
HDX-MS time courses for peptides with significant differences. **A**. Selected Akt1 peptides that showed decreases and increases in exchange. Error is shown as standard deviation (n=3). **B**. Selected Akt1 3P peptides that showed decreases and increases in exchange. Error is shown as standard deviation (n=3). **C**. Selected Akt1 3P peptides that showed decreases and increases in exchange. Error is shown as standard deviation (n=3). The location of these peptides is indicated in the title, with the sequence indicated. Full HDX-MS data for all experiments is in the source data.

**Figure S4:**
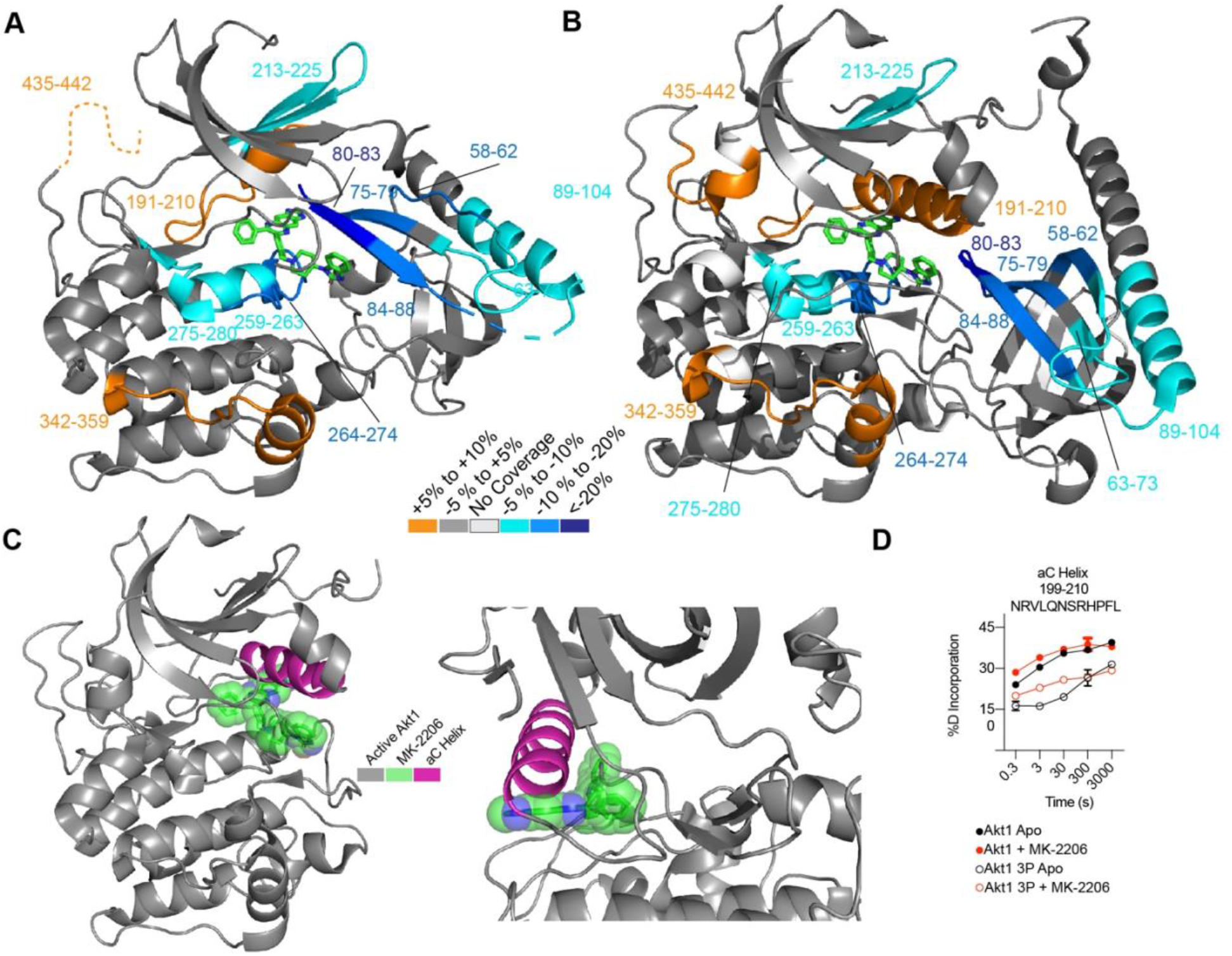
Mapping changes upon MK-2206 binding on different structural models, and putative clash between activated αC helix with MK-2206. **A+B**. Peptides showing significant deuterium exchange differences (>4.5%, 0.45 Da, and p<0.01in an unpaired two-tailed T-test) between Akt1 3P apo vs. Akt1 3P + MK-2206. Significant differences in deuterium exchange are mapped according to the legend on the structure of MK-2206 bound to Akt1 (panel **A**, PDB: 3O96) and a model composed of the kinase domain from phosphorylated Akt1 (4EKK), the PH domain from inactive Akt1 (7APJ), and the MK-2206 from (3096) (panel **B**). **C**. Model of fully phosphorylated Akt1 (4EKK), aligned to MK-2206. Fully phosphorylated Akt1 kinase domain (PDB: 4EKK) aligned to Akt1 in complex with MK-2206 (PDB: 3O96). Zoom indicates the putative clashes between the αC helix of phosphorylated Akt1 with MK-2206. **D**. αC helix peptide that showed significant protection between unphosphorylated and phosphorylated Akt1 (due to formation of αC helix), and increased exchange in the presence of MK-2206. Error is shown as standard deviation (n=3).

**Supplementary Table 1.**
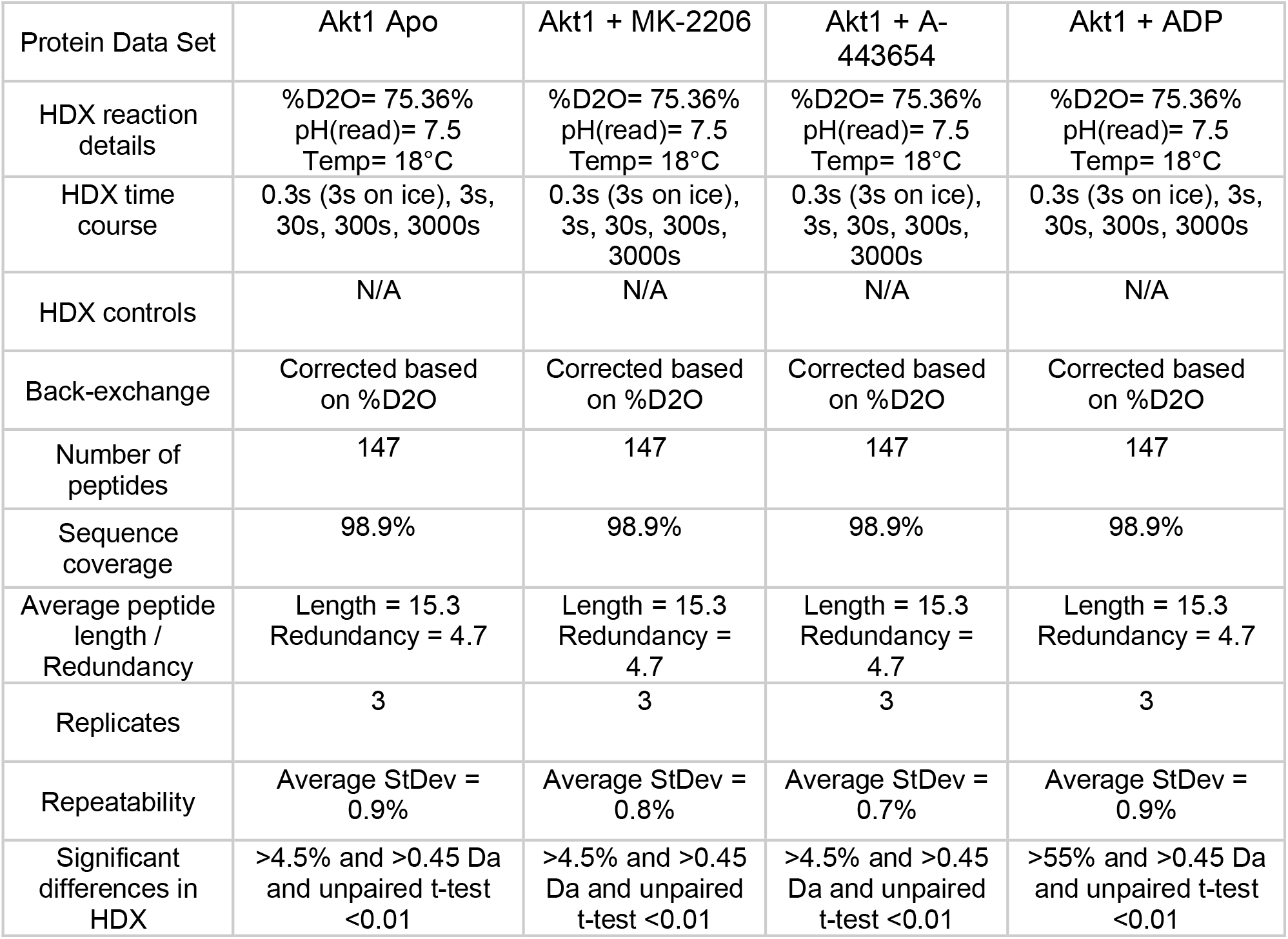
Akt1 HDX Data Summary.

**Supplementary Table 2.** Akt1 3P HDX Data Summary.

| Protein Data Set | Akt1 3P Apo | Akt1 3P + MK-2206 | Akt1 3P + A-443654 | Akt1 3P + 5% PIP3 Vesicles | Akt1 3P + 5% PIP3 Vesicles + MK-2206 | Akt1 3P + 5% PIP3 Vesicles + A-443654 |
| --- | --- | --- | --- | --- | --- | --- |
| HDX reaction details | %D2O= 66%<br>pH(read)= 7.5 | %D2O= 66%<br>pH(read)= 7.5 | %D2O= 66%<br>pH(read)= 7.5 | %D2O= 66%<br>pH(read)= 7.5 | %D2O= 66%<br>pH(read)= 7.5 | %D2O= 66%<br>pH(read)= 7.5 |
| HDX time course | 0.3s (3s on ice), 3s, 30s, 300s, 3000s | 0.3s (3s on ice), 3s, 30s, 300s, 3000s | 0.3s (3s on ice), 3s, 30s, 300s, 3000s | 3s, 30s, 300s, 3000s | 3s, 30s, 300s, 3000s | 3s, 30s, 300s, 3000s |
| HDX controls | N/A | N/A | N/A | N/A | N/A | N/A |
| Back-exchange | Corrected based on %D2O | Corrected based on %D2O | Corrected based on %D2O | Corrected based on %D2O | Corrected based on %D2O | Corrected based on %D2O |
| Number of peptides | 143 | 143 | 143 | 143 | 143 | 143 |
| Sequence coverage | 98.9% | 98.9% | 98.9% | 98.9% | 98.9% | 98.9% |
| Average peptide length / Redundancy | Length = 14.9<br>Redundancy = 4.5 | Length = 14.9<br>Redundancy = 4.5 | Length = 14.9<br>Redundancy = 4.5 | Length = 14.9<br>Redundancy = 4.5 | Length = 14.9<br>Redundancy = 4.5 | Length = 14.9<br>Redundancy = 4.5 |
| Replicates | 3 | 3 | 3 | 3 | 3 | 3 |
| Repeatability | Average StDev = 0.7% | Average StDev = 0.7% | Average StDev = 0.8% | Average StDev = 1.2% | Average StDev = 0.9% | Average StDev = 0.9% |
| Significant differences in HDX | >4.5% and >0.45 Da and unpaired t-test <0.01 | >4.5% and >0.45 Da and unpaired t-test <0.01 | >4.5% and >0.45 Da and unpaired t-test <0.01 | >4.5% and >0.45 Da and unpaired t-test <0.01 | >4.5% and >0.5 Da and unpaired t-test <0.01 | >4.5% and >0.5 Da and unpaired t-test <0.01 |

